# Multi-pulse transcranial magnetic stimulation of human motor cortex produces short-latency corticomotor facilitation via two distinct mechanisms

**DOI:** 10.1101/2022.02.19.481138

**Authors:** Janine Kesselheim, Mitsuaki Takemi, Lasse Christiansen, Anke Ninija Karabanov, Hartwig Roman Siebner

## Abstract

**Background:** Single-pulse transcranial magnetic stimulation of the precentral hand representation (M1_HAND_) can elicit indirect waves in the corticospinal tract at a periodicity of ~660 Hz, called indirect or I-waves. These synchronized descending volleys are produced by transsynaptic excitation of fast-conducting monosynaptic corticospinal axons in M1-HAND. Paired-pulse TMS can induce short-interval intracortical facilitation (SICF) of motor evoked potentials (MEPs) at inter-pulse intervals that match I-wave periodicity.

**Objective:** To examine whether short-latency corticospinal facilitation engages additional mechanisms independently of I-wave periodicity.

**Methods:** In 19 volunteers, one to four biphasic TMS pulses were applied to left M1-HAND with interpulse interval was adjusted to the first peak or first trough of the individual SICF curve. TMS was applied at different intensities to probe the intensity-response relationship.

**Results:** Pairs, triplets, or quadruplets at individual peak-latency facilitated MEP amplitudes across a wide range of TMS intensities compared to single pulses. Multi-pulse TMS_HAND_ at individual trough-latency also produced a consistent facilitation of MEP amplitude. Short-latency facilitation at trough-latency was less pronounced than short-latency facilitation at peak-latency, but the relative difference in facilitation decreased with increasing stimulus intensity. Increasing the number of pulses from two to four pulses had only a modest effect on MEP facilitation.

**Conclusion:** Two mechanisms underly short-latency corticomotor facilitation caused by biphasic multi-pulse TMS. An intracortical mechanism is related to I-wave periodicity and engages fast-conducting direct projections to spinal motoneurons. A second corticospinal mechanism does not rely on I-wave rhythmicity and may be mediated by slower conducting indirect pyramidal tract projections from M1-HAND to spinal interneurons. The latter mechanism deserves more attention in TMS studies of the corticomotor system.

## Introduction

Transcranial magnetic stimulation of the precentral motor hand representation (M1_HAND_) has been widely used to study the physiology of the human corticomotor system [1, 2]. A single TMS pulse can elicit multiple short-latency volleys in the descending pyramidal tract [3]. At intensities that are slightly above the corticomotor threshold, TMS of M1_HAND_ elicits indirect waves (I-waves), which display a periodicity of approximately 660 Hz [3]. It is widely accepted that these I-waves are caused by transsynaptic (indirect) excitation of layer-5 pyramidal neurons in M1-HAND that make direct synaptic connections with the spinal motoneurons [4–11]. The number of I-waves and their amplitude increase with the intensity of TMS, while a directly evoked descending volley (D-wave) with an even shorter latency first emerges at high stimulus intensities [12, 13].

While the descending waves can only be studied invasively via epidural recordings [14, 15], the periodicity of I-waves can be studied non-invasively using paired-pulse TMS of M1_HAND_ [16, 17]. When paired-pulse TMS is applied through the same coil at a short inter-pulse interval (IPI), the amplitude of the motor evoked potentials (MEPs) are facilitated in the contralateral hand, showing distinct peaks of short-interval intracortical facilitation (SICF) of the muscle response at IPIs of 1.0-1.5, 3.0 and 4.5 ms [16, 17]. The similarity between the periodicity of the SICF curve and descending I-waves led to the notion that the paired-pulse TMS paradigm used to probe SICF acts on the intracortical circuitry that generate I-waves.

Since SICF is strongest at an IPI of 1.5 ms, paired-pulse repetitive TMS protocols (pp-rTMS) at an IPI of 1.5 ms have been used to boost the facilitatory after-effects of repetitive TMS on corticomotor excitability. Other studies used high-frequency bursts at IPIs within the SICF range to stimulate M1-HAND and showed that four-pulse bursts (quadruplets) at an IPI of 1.5 ms produces lasting bidirectional effects on corticospinal excitability [18–20]. The direction of the excitability change (i.e., facilitation or suppression) after quadruple TMS critically depended on the repetition rate of the quadruplets [18–20]. Together, these studies have raised considerable interest in multi-pulse TMS at an ISI that corresponds to I-wave periodicity in interventional studies that are aiming at inducing plasticity in the human corticomotor system.

Multi-pulse TMS at short IPIs may not only facilitate those circuits in M1-HAND that underpin SICF of fast-conducting direct corticospinal projections, but also cause substantial short-latency facilitation of motor evoked responses via other mechanisms. This hypothesis is supported by a study in macaques, in which excitatory postsynaptic potentials (EPSPs) were recorded from cervical motoneurons after intracortical stimulation of the motor cortex[21]. In that study, EPSPs were evoked by single-pulse, paired-pulse or triple pulses using a short IPI of 3 ms. The recordings revealed that a substantial portion of the cortically evoked EPSPs were caused by excitation of polysynaptic corticospinal projection. These polysynaptically generated EPSPs, but not the EPSPs generated via monosynaptic corticomotoneuronal projections, increased in amplitude and shortened response onset latency with multi-pulse stimulation relative to single-pulse stimulation.

Motivated by this intriguing finding in macaques, we applied single-, paired-, triple-, and quadruple-pulse TMS to left M1-HAND across a wide range of stimulus intensities to study short-latency corticomotor facilitation in healthy human volunteers. We hypothesized that short-latency facilitation also engages corticospinal mechanisms that does not rely on I-wave periodicity. Since TMS triplets with IPIs of 1.5 ms have been shown to facilitate MEPs to a larger extent than doublets [22], we expected an increase in the efficacy of multi-pulse TMS_HAND_ with increasing number of stimuli due to a more effective repetitive activation of the cortical or spinal interneuron pools contributing to short-latency multi-pulse facilitation.

## Material and methods

All experimental procedures were in accordance with the latest revision of the Declaration of Helsinki and approved by the local Ethics Committee (De Videnskabsetiske Komiteer, Region Hovedstaden, Journal-Nr. H-15017238). All participants gave written informed consent.

### Experimental design

Experimental procedures consisted of a main experiment and two follow-up experiments (see figure 1), conducted at least 48 h apart to avoid carry-over effects. In the main and first followup experiment, we applied single-, paired-, triple- and quadruple-pulse TMS_HAND_ at two IPIs. The second follow-up experiment applied single- and paired-pulse TMS_HAND_ at four different IPIs.

**Fig. 1.**
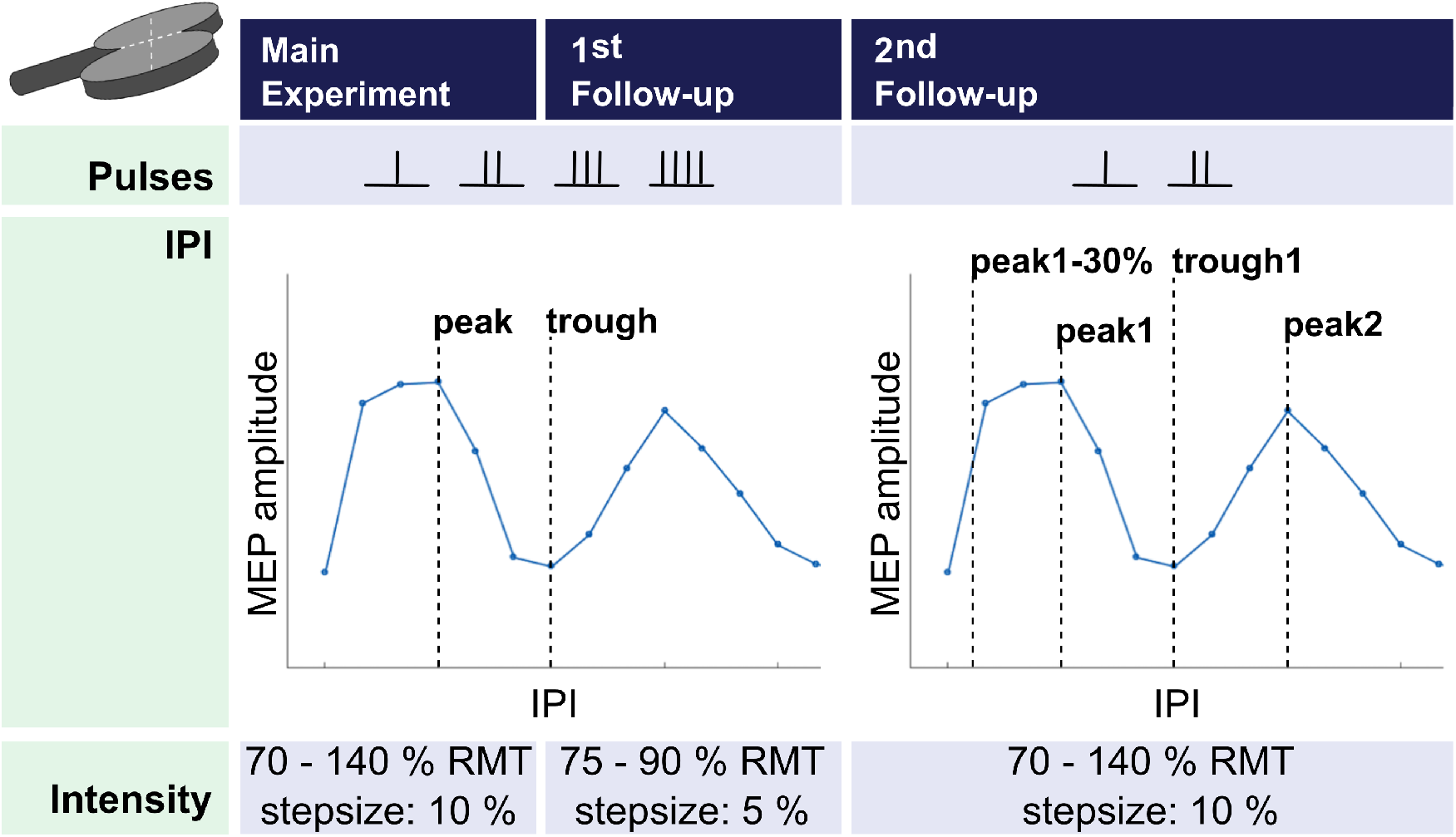
Schematic overview of the three experiments. The main and first follow-up experiment applied single-, paired-, triple- and quadruple-pulse TMS_HAND_ in two different inter-pulse intervals (IPIs), while the second follow-up applied only single- and paired-pulse TMS_HAND_ in four different IPIs. TMS intensities along with their stepsizes are represented in the last row.

### Subjects

Twenty-one participants (eight men; age range: 19-35 years, mean age: 25 years) took part in the main experiment. 11 subjects also participated in the first and 17 in the second follow-up experiment. Participants were right-handed according to the Edinburgh Handedness Inventory (mean laterality quotient: 93.1, SD:14.0) apart from one participant being ambidextrous with a laterality quotient of 43. Exclusion criteria were pregnancy, intake of medication acting on the central nervous system, implanted medical devices, history of neurological disorders and claustrophobia.

#### Transcranial magnetic stimulation

Prior to the TMS session, each participant underwent T1-weighted magnetic resonance imaging (MRI) of the whole brain (Philips Achieva 3T MRI scanner). The T1 was used for frameless neuronavigation of the TMS coil (Localite, Sankt Augustin, Germany) to secure coil position over the motor hotspot of the right first dorsal interosseus (FDI) muscle. TMS was applied with a standard MC-B70 figure-of-eight coil attached to a MagPro X100 Option stimulator (MagVenture, Farum, Denmark). The flat surface of the coil touched the scalp with the handle directed posteriorly in a 45° angle to the sagittal midline. All pulses delivered during the main and follow-up experiments were biphasic, where the second phase induced a posterior-to-anterior (P-A) current direction. We determined the motor hotspot, defined as the coil position, at which a single TMS pulse elicited the largest MEPs in the right FDI muscle. Pulses were delivered with an inter-trial interval of five seconds with a 25% jitter. Individual RMT and stimulation intensity needed to elicit MEPs with a peak-to-peak amplitude of ~1mV (TMS_1mV_) were determined using adaptive parameter estimation by sequential testing and maximum likelihood regression [23].

#### Electromyographic recordings

Participants were resting in a comfortable chair with their eyes open. MEPs were recorded with self-adhesive surface Ag/AgCl electrodes (Ambu Neuroline 700, Columbia, USA) mounted on thoroughly cleaned skin above the right FDI muscle in a muscle-tendon montage. EMG signals were sampled at 5 kHz, band-pass filtered (5-2000 Hz) and amplified (x1000) using eight-channel DC amplifier (1201 micro Mk-II unit, Digitimer, Cambridge Electronic Design) and Signal software version 4.11 (Cambridge Electronic Design, Cambridge, UK).

### Main experiment

Using SICF-adjusted multi-pulse TMS, we assessed the influence of number of pulses and IPI on the stimulus-response curve. The IPIs were personalized based on the individual SICF curve.

#### Short-interval cortical facilitation (SICF)

To estimate the individual peak-latencies, we constructed SICF-curves illustrating the MEP facilitation from paired pulses with IPIs from 1.1 to 2.7ms in steps of 0.2ms (Fig. 1). The intensity of the first pulse was set to TMS_1mV_ and the second pulse to 90% RMT. Ten paired pulses per IPI and ten single pulses were delivered in a randomized manner. The latencies of the first peak and trough of the SICF curve were identified as the IPIs at which the highest and lowest MEP amplitudes occurred (Fig. 1).

#### Stimulus-response curve obtained with SICF-adjusted multi-pulse TMS

Biphasic TMS pulses were applied either as single pulses or pairs, triplets, or quadruplets. The IPI within the bursts was constant and adjusted to either the individual latency of the first peak (peak1-latency) or the trough between the first and second peak (trough1-latency). Ten MEPs per pulse number and IPI were recorded at eight different TMS intensities (70, 80, 90, 100, 110, 120, 130, 140% RMT). The stimulation intensity was identical for all pulses within a burst. The order of TMS intensities was randomized and counter-balanced across participants. For each TMS intensity, the seven different TMS conditions (single-pulse and pairs, triplets, and quadruplets at peak- or trough-latency) were tested in a randomized order.

### First follow-up experiment: SICF-adjusted TMS at sub-motor threshold intensities

Since the cumulative effects of multiple pulses may be affected by ceiling effect at higher intensities, we conducted a follow-up that focused on a low TMS intensity range from 75 to 90% RMT (Fig. 1). We used increments of 5% RMT to probe the facilitatory effects in the low-intensity range in greater detail. Otherwise, the experimental procedures were identical to the main experiment.

### Second follow-up experiment: Impact of temporal proximity

To exclude the possibility that higher facilitation at individual peak compared to through IPIs due to a shorter IPI, we conducted a third experiment including an IPI 30% shorter than the first peak (peak1-30%-latency) and an IPI corresponding to the second SICF peak (peak2-latency) (Fig. 1).

We again assessed the stimulus-response curve, covering the same intensity range as in the first experiment. The design included 40 conditions: five pulse conditions, namely single-pulse and paired-pulse stimulation at four different IPIs (peak1-30%-latency, peak1-latency, trough1-latency, peak2-latency) at eight TMS intensities. Conditions were pseudo-randomized with 10 trials per condition.

### Data analysis

All MEP trials were visually inspected for voluntary muscle activity and trials showing background EMG activity during the 50-ms period before stimulation were discarded. We constructed stimulus-response curves for each combination of pulse number and IPI, showing the increase in mean MEP amplitude with increasing stimulus intensity. Mean MEP amplitudes were computed from MEP amplitudes extracted trial-by-trial. We also calculated the difference in mean MEP amplitude evoked by pairs, triplets, and quadruplets relative to single-pulse TMS and constructed normalized stimulus-response curves.

The averaged and normalized MEP amplitudes for each subject and experimental condition were entered in a full-factorial analysis of variance (ANOVA). MEP amplitudes were log-transformed prior to performing statistical analysis to ensure normal distribution of the data. We computed three-way ANOVAs to analyse the normalized MEP amplitudes recorded in the main experiment. The ANOVAs treated the factors number of pulses (pairs, triplets, and quadruplets), IPI (peak1-latency and trough1-latency) and TMS intensity (70–140% RMT) as within-subject factors.

The same ANOVA model was specified to analyse normalized MEP amplitudes in the first follow-up with the within-subject factors number of pulses (pairs, triplets, and quadruplets), IPI (peak1-latency and trough1-latency) and TMS intensity (75–90 % RMT). In separate follow-up ANOVAs, we assessed differences in normalized MEP amplitudes across conditions for each TMS intensity level separately. For the main experiment and the first follow-up, the ANOVA model included the within-subject factors pulse number (pairs, triplets, and quadruplets) and IPI (peak1 and trough1).

Mean MEP amplitudes recorded in the second follow-up experiment were analysed using two separate two-way ANOVAs to directly compare the conditions peak1-30% and peak1 and the conditions trough1 and peak2, respectively. The two ANOVA models treated the factors IPI (peak1-30%-latency, peak1-latency, trough1-latency, peak2-latency) and TMS intensity (70– 140% RMT) as within-subject factors. Again, in separate follow-up ANOVAs, we assessed differences in mean MEP amplitudes across conditions for each TMS intensity level separately with the within-subject factor IPI. Bonferroni corrections for multiple comparisons were applied. All ANOVAs used Mauchly’s Test for Sphericity and Greenhouse-Geisser Sphericity Corrections and were conducted using the SPSS 25 software package (IBM Corp., New York, USA). Significance threshold was set to 0.05.

## Results

Two volunteers in the first experiment dropped out due to perceived discomfort during TMS. None of the remaining 19 participants (seven men; age range: 19-35 years, mean age: 25 years) reported discomfort or side effects. In one participant, we failed to record the data at 100 % RMT due to a technical error in the second follow-up experiment, resulting in a total of 16 participants for this TMS intensity. All data are available in “Open Science Framework” at https://osf.io/acges/ (DOI 10.17605/OSF.IO/ACGES, Kesselheim et al., 2020).

### Short-interval intracortical facilitation

Figure 2 illustrates the group SICF data. While all participants showed a clear peak-1 and peak-2 separated by a trough (trough-1), peak1- and trough1-latencies varied across individuals. This resulted in an overlap of the maximally observed peak1-latency (1.7 ms) and the minimally observed trough1-latency (1.7 ms) across participants and experiments, providing a post-hoc justification for personalizing the IPIs according to the individual SICF curve.

**Fig. 2.**
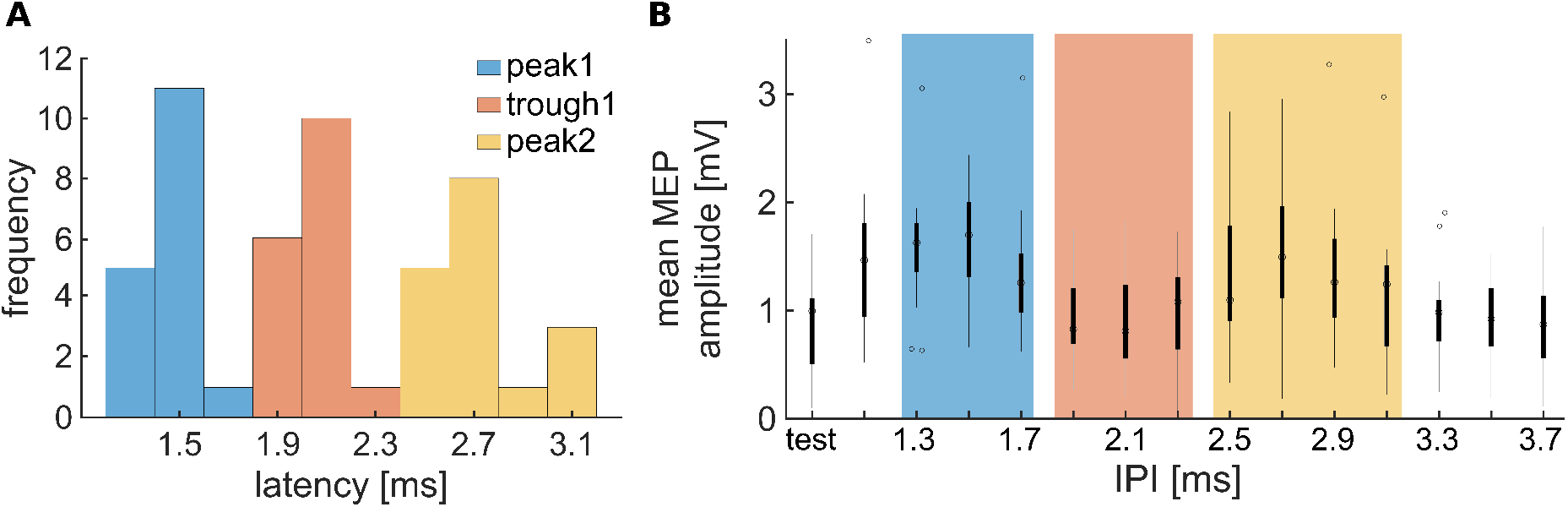
SICF latency distribution and profile. **(A)** distribution of the different peak- and trough-latencies obtained by SICF in 17 healthy participants in the second follow-up experiment, **(B)** grand average SICF-profile represented in boxplots of the same experiment; boxes show the median MEP amplitude (central dot) and the 25th (bottom edge) and 75th percentile (top edge), the whiskers extend to the highest and lowest mean MEP amplitude included in the analysis, while outliers are plotted individually as circles. In SICF, the test pulse was a single pulse, thereafter pairs were applied in stepsizes of 0.2 ms IPI starting at 1.1 ms.

### Main experiment

Multi-pulse TMS consistently caused short-latency facilitation of the normalized MEP amplitudes (Fig.3). Separate two-way ANOVAs showed that this was the case for both, multipulse TMS_HAND_ at peak1-latency and trough1-latency. Multi-pulse TMS_HAND_ at peak1-latency enhanced the mean MEP amplitude relative to single-pulse TMS. This facilitatory effect increased with pulse number (F (2.34,42.1) = 176, p = 3.5e-22) and stimulus intensity (F (2.51,45.1) = 161, p = 6.9e-23). The ANOVA also showed a significant interaction between pulse number and stimulus intensity (F (6.7,121) = 12.5, p = 1.2e-11). Multi-pulse TMS_HAND_ at trough1-latency also produced short-latency facilitation (Fig.3A). There was a significant main effect of pulse number (F (2.32,41.9) = 44.8, p = 7.3e-12) and stimulus intensity (F (2.65,47.7) = 205, p = 2.7e-26) as well as a significant interaction between pulse number and stimulus intensity (F (6.90,124) = 2.99, p = 0.0065).

**Fig. 3.**
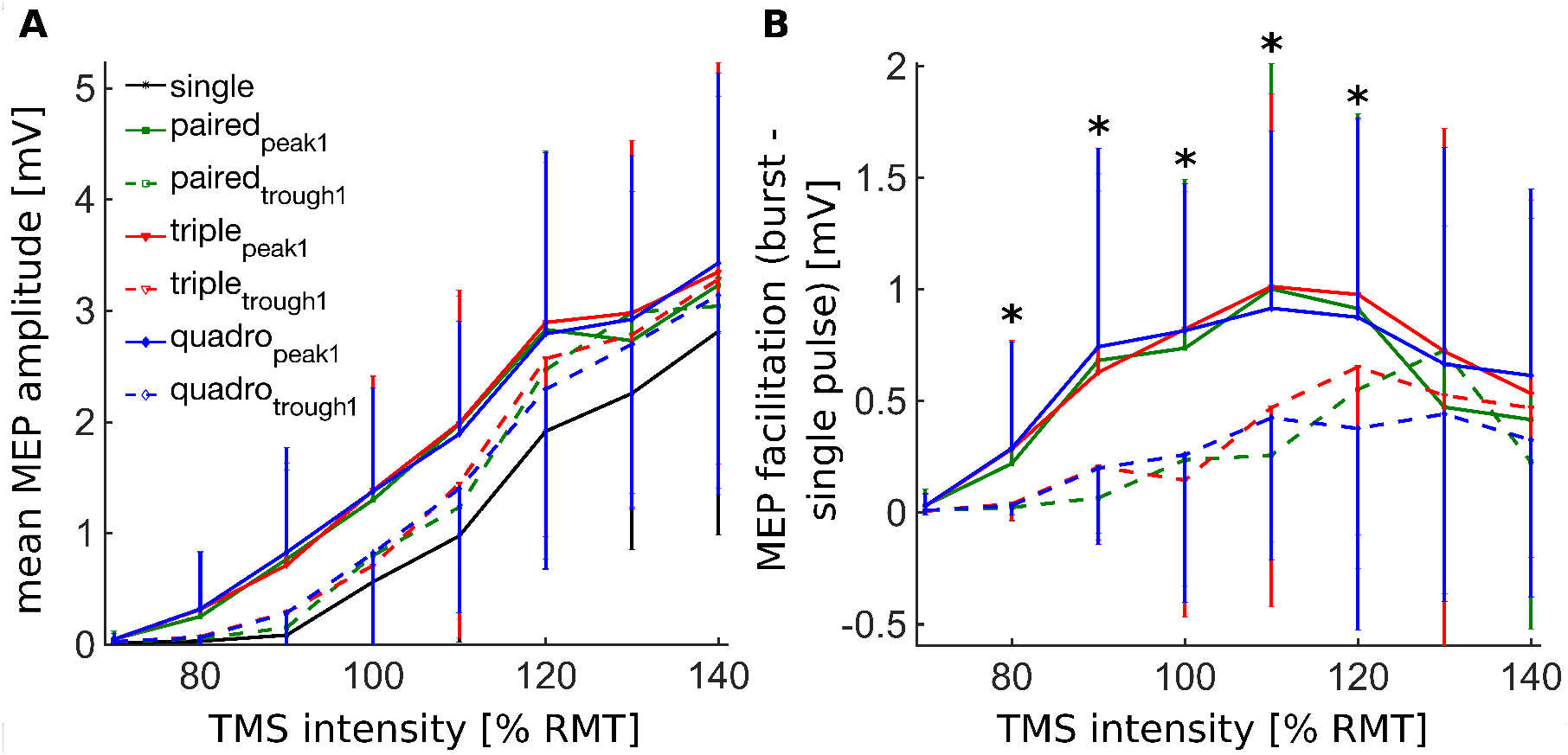
Peak and trough multi-pulse facilitation. Mean group data showing the increase in mean MEP amplitudes with TMS intensity (main experiment, *n* = 19). Error bars represent the standard deviation of the mean for each configuration. *Legend:* single - single-pulse TMS_HAND_, paired-paired-pulse TMS_HAND_, triple - triple-pulse TMS_HAND_, quadro - quadruple-pulse TMS_HAND_, *indices:* IPI at the latency of the first peak in SICF, trough - IPI at the latency of the trough between the first two peaks in SICF. (**A**) absolute mean MEP amplitude for TMS intensities from 70 to 140% resting motor threshold (RMT), (**B**) MEP facilitation is represented as absolute difference between mean MEP amplitudes of multi-pulse and single-pulse TMS_HAND_. Asterisks mark significant effects of IPI on MEP facilitation (80% RMT: *p* = 0.0023; 90% RMT: *p* < 0.0001; 100% RMT: *p* < 0.0001; 110% RMT: *p* = 0.0003; 120% RMT: *p* = 0.0051; two-way ANOVA, Bonferroni corrected). Both trough1- and peak1-adjusted multi-pulse TMS_HAND_ show no MEP facilitation at 70% RMT. This is because neither multi-pulse nor single-pulse TMS_HAND_ was able to elicit MEPs at an intensity as low as 70% RMT.

While multi-pulse TMS_HAND_ at peak1-latency and trough1-latency consistently produced shortlatency facilitation, there were also notable differences between the two conditions in terms of short-latency MEP facilitation (Fig.3B). This was evidenced by a three-way ANOVA of multipulse MEP amplitudes normalized to single pulse MEP amplitude, including the factors ‘IPI’, ‘TMS intensity’ and ‘pulse number’. Like the two-way ANOVAs, three-way ANOVA showed significant main effects of TMS intensity (*F* (3.32,59.7) = 13.6, *p* = 3.04e-7) and pulse number (*F* (1.71,30.9) = 11.6; *p* = 3.20e-4). In addition, the three-way ANOVA revealed a main effect of IPI (*F* (1,18) = 119, *p* = 2.24e-9) and a significant interaction between TMS intensity and IPI (*F* (3.14,56.6) = 14.83, *p* = 2.07e-7). This interaction can be attributed to a relatively stronger short-latency MEP facilitation of multi-pulse TMS_HAND_ at peak1-latency in the lower TMS intensity range. Post-hoc analyses showed that multi-pulse TMS_HAND_ at peak1-latency produced a stronger facilitatory effect at TMS intensities from 80 to 120% RMT than multi-pulse TMS_HAND_ at trough1-latency (all: *F* (1,18) = 20.1, all *p* = 0.0052). The three-way ANOVA did not reveal additional interactions among the three factors.

Together, the results demonstrate that personalized multi-pulse TMS at individual peak1-latency or through1-latency excite the corticospinal system more effectively than single-pulse TMS. While the strength of short-latency corticospinal facilitation becomes more and more comparable with increasing stimulus intensity, the magnitude of short-latency corticospinal facilitation at individual peak1-latency is larger than short-latency at individual trough1-latency within the lower intensity range of the stimulus-response curve.

### First follow-up experiment

In this experiment, we probed short-latency corticospinal facilitation at sub-motor threshold intensities using smaller increments in intensity (5% of individual RMT). Multi-pulse TMS_HAND_ adjusted to individual peak1-latency induced a stronger MEP facilitation relative to single-pulse TMS_HAND_ and multi-pulse TMS_HAND_ adjusted to individual trough1-latency (Fig. 4). This was reflected by the ANOVA, showing a main effect of IPI (*F* (1,10) = 19.0, *p* = 0.0014). The stronger facilitation with multi-pulse TMS at peak1-latency first emerged at TMS intensities close to the RMT. Accordingly, the ANOVA showed a main effect of stimulus intensity (*F* (1.32,13.2) = 7.38, *p* = 0.0126) and a significant interaction between IPI and stimulus intensity (*F* (1.42,14.2) = 16.4, *p* = 5.0e-4). Post-hoc testing showed that a significant facilitatory effect for peak1-latency adjusted multi-pulse TMS on MEP amplitude was first observed at intensities corresponding to 85 and 90% RMT (85%: *F* (1,10) = 12.0, *p* = 0.0243; 90%: *F* (1,10) = 57.7, *p* = 7.6e-5), but not at lower TMS intensities (75%: *F* (1,10) = 6.66,*p* = 0.110; 80%: *F* (1,10) = 3.80, *p* = 0.320).

**Fig. 4.**
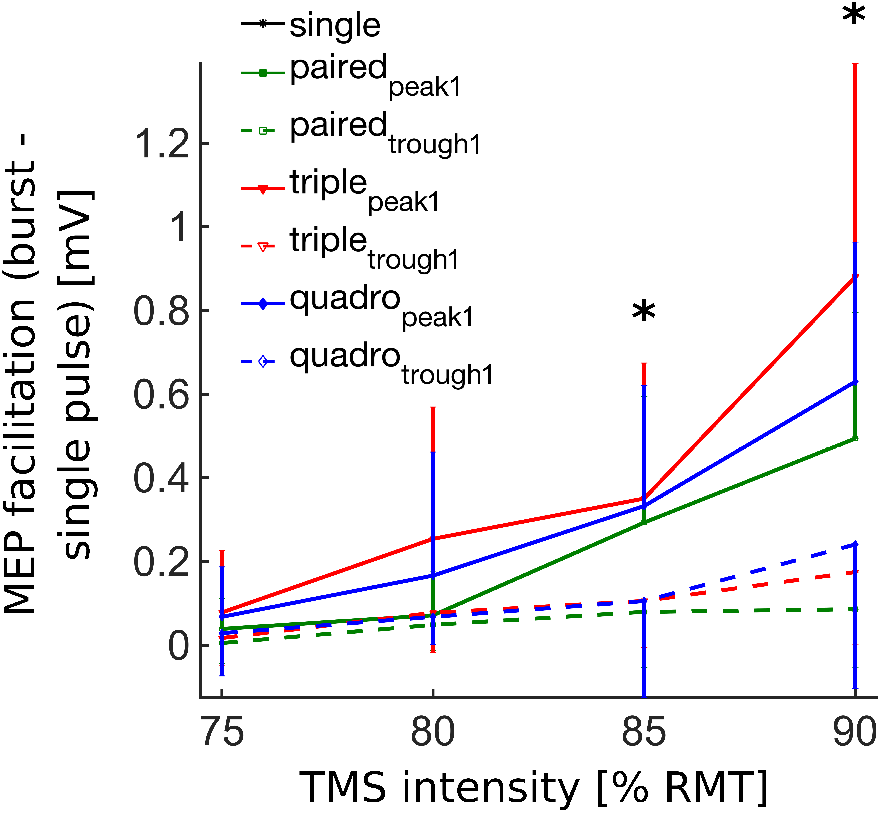
Multi-pulse facilitation at subthreshold intensities. Mean group data obtained from the first follow-up experiment (*n* = 11); MEP facilitation represented as absolute difference between mean MEP amplitudes of multi-pulse and singlepulse TMS_HAND_. Error bars represent the standard deviation of the mean for each configuration. Asterisks mark significant effects of IPI on MEP facilitation (85% RMT: *p* = 0.0243; 90% RMT: *p* < 0.0001; two-way ANOVA, Bonferroni corrected).

At the sub-motor threshold intensities tested, the number of TMS_HAND_ pulses contributed to MEP facilitation. The ANOVA showed a main effect of pulse number (*F* (1.24,12.4) = 13.5, *p* = 0.0020). This main effect was caused by higher MEP amplitudes with a higher number of pulses (Fig. 4). This facilitatory effect was independent of the IPI and stimulus intensity, reflected by a lack of interaction between IPI and pulse number (*F* (1.15,11.5) = 0.39, *p* = 0.574) or TMS intensity and pulse number (*F* (4.34,43.4) = 2.06, *p* = 0.0971).

### Second follow-up experiment

In this experiment, we tested how much temporal closeness of consecutive TMS pulses affect MEP amplitude as opposed to SICF periodicity (Fig. 5). In this experiment, we compared MEP amplitude elicited by paired-pulse TMS_HAND_ at four IPIs, including an IPI that was 30% shorter than individual peak1-latency (peak1-30%-latency) and an IPI corresponding to the second SICF peak (peak2-latency).

**Fig 5.**
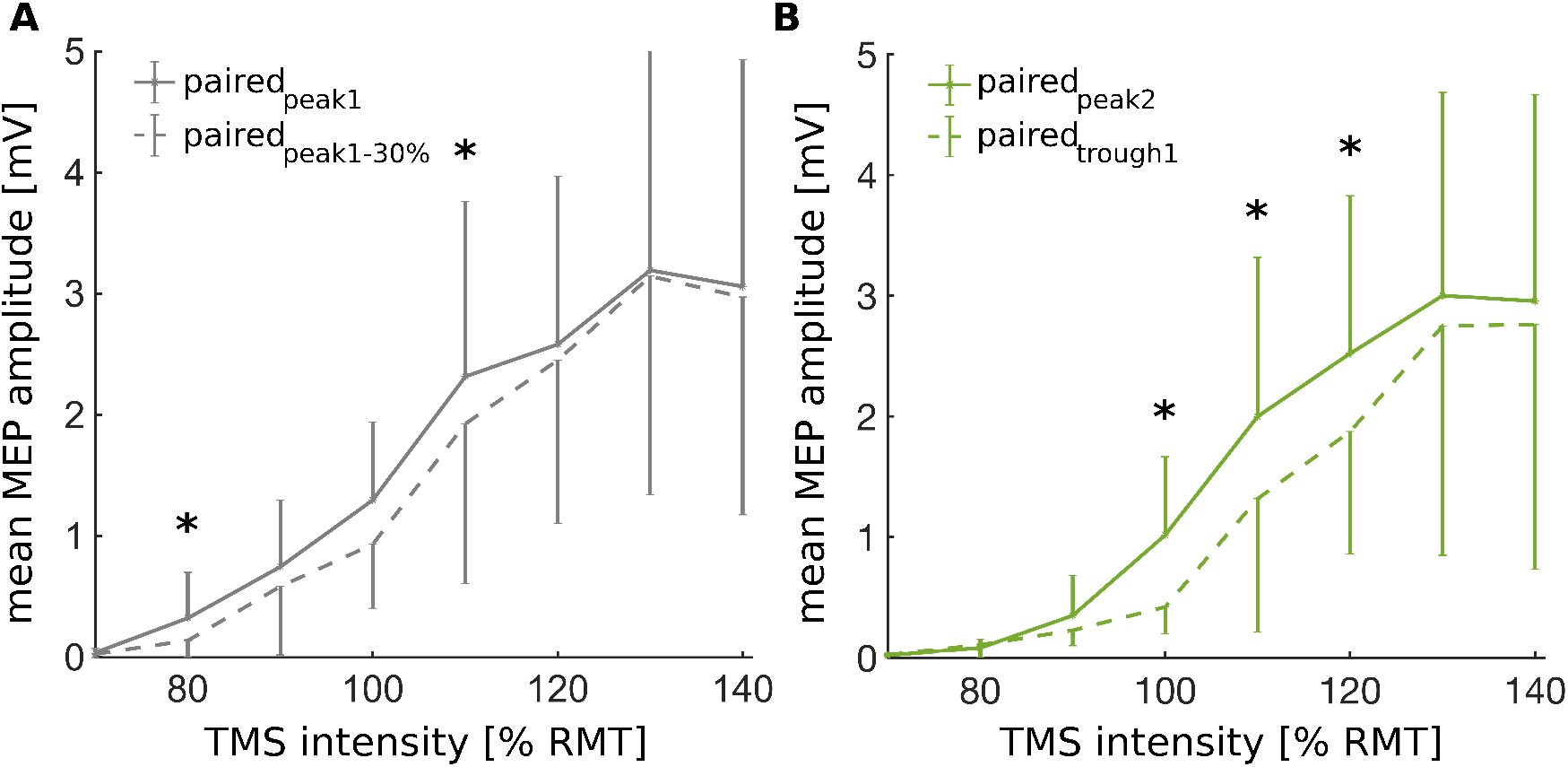
Effect of temporal proximity on paired-pulse MEP amplitude. Mean group data of the second follow-up experiment (*n* = 17) represented as absolute mean MEP amplitudes of paired-pulse TMS_HAND_ at TMS intensities from 70 to 140% RMT. **(A)** mean MEP amplitudes at IPIs corresponding to the first SICF peak (peak1, solid line) and 30% less than the first peak (peak1-30%, dashed line) **(B)** mean MEP amplitudes at IPIs corresponding to the trough between the first and second peak (trough1, dashed line) and the second SICF periodicity peak (peak2, solid line). Error bars represent the standard deviation of the mean for each configuration. Asterisks mark significant effects of IPI on MEP amplitude (peak1 vs. peak1-30%: 80% RMT: *p* = 0.016; 110% RMT: *p* = 0.0082; peak2 vs trough1: 100% RMT: *p* = 0.011; 110% RMT: *p* = 1.4e-4; 120% RMT: *p* < 8e-6; one-way ANOVAs, Bonferroni corrected).

Confirming the added response magnitude due to SICF periodicity, paired pulses adjusted to peak1-latency consistently elicited larger MEPs than paired pulses at peak1-30%-latency (Fig.5a). Likewise, paired pulses at an IPI adjusted to peak2-latency elicited larger MEPs than paired pulses at individual trough-1 latency (Fig.5b). Comparing paired-pulse TMS at peak1-latency and peak1-30%-latency, two-way ANOVA showed a significant effect of IPI (*F* (1,15) = 12.9, *p* = 0.0027), stimulus intensity (*F* (2.36,35.4) = 225.9, p = 7.9e-22) and an interaction between IPI and intensity (*F* (3.14,47.1) = 4.28, *p* = 0.0086). Follow-up ANOVAs demonstrated that paired-pulse TMS at peak1-latency produced more MEP facilitation than paired-pulse TMS at peak1-30%-latency, when stimulus intensity was set at 80% RMT (*F* (1,16) = 7.1, *p* = 0.0158) and 110% RMT (*F* (1,16) =16.0, *p* = 0.0082).

MEP amplitudes evoked by a paired-pulse TMS at peak2-latency evoked larger MEP amplitudes than paired-pulse TMS at trough1-latency. Again, the two-way ANOVA showed a significant effect of IPI (*F* (1,15) = 11.2, *p* = 0.0044), stimulus intensity (*F* (3.18,47.8) = 292.5, *p* = 3.1e-31) and an interaction between IPI and intensity (*F* (2.58,38.7) = 5. 78, *p* = 0.0042). The interaction can be attributed to a larger facilitatory effect of paired-pulse TMS at peak2-latency at intermediate stimulus intensities (Fig. 5B). Post-hoc testing showed stronger MEP facilitation for peak2-latency adjusted paired-pulse TMS at 100% RMT (*F* (1,15) = 15.5, *p* = 0.011), 110% RMT (*F* (1,16) = 36.7, *p* = 1.4e-4), and 120% RMT (*F* (1,16) = 56.7, *p* = 8.0e-6).

In summary, the facilitatory of paired-pulse TMS at SICF periodicity cannot solely be attributed to temporal proximity of the two pulses since both analyses confirmed that longer IPIs adjusted to SICF peaks are more efficient in eliciting corticomotor responses as compared to shorter IPIs not matching SICF peaks.

## Discussion

We found that multi-pulse TMS at ultra-high repetition rate facilitates corticomotor excitability via two mechanisms. We replicated the well-known short-latency facilitation when the interval between consecutive pulses was adjusted to individual latency of the first peak in the SICF curve, corresponding to a repetition rate of ~660 Hz that matches I-wave periodicity. Expanding the existing knowledge, we still found a consistent facilitatory effect, when the interval between consecutive pulse was adjusted to the individual latency of the first trough in the SICF curve. This observation demonstrates the existence of short-latency cortico-motor facilitation that is independent of I-wave periodicity.

### Effects of multi-pulse TMS at peak1-latency

When we adjusted the IPI to the individual latency of the first SICF peak, multi-pulse TMS_HAND_ elicited larger MEPs than multi-pulse TMS_HAND_ with an IPI adjusted to trough1-latency. Hence, the short-latency facilitatory effect was strongest, when TMS_HAND_ exploited I-wave periodicity. This observation confirms our hypothesis that adjusting the IPI to individual I-wave periodicity increased the efficacy of stimulation-induced excitation of the fast-conducting corticospinal output neurons in M1_HAND_. The results obtained in the second control experiment speaks against the possibility that the higher efficacy of multi-pulse TMS_HAND_ at individual SICF periodicity was merely caused by a shorter IPI (i.e. a higher stimulation rate).

Due to the striking similarities between the temporal profiles of SICF and invasive I-wave recordings, it has been suggested that SICF reflects facilitation of intracortical circuits that create the multiple I-waves in M1_HAND_ [16, 17]. As already mentioned in the introduction, I-waves reflect synchronized activity in fast-conducting corticospinal neurons (i.e., descending waves). Whereas there is an agreement that the intrinsic properties of the fast-conducting corticospinal neurons are critical (but not sufficient [24]) to generate I-waves, the precise physiological mechanisms are still a matter of debate [3, 25]. The fast-conducting corticospinal projections have very short refractory periods enabling high-frequency firing [26]. Such firing have been proposed to result from repetitive transsynaptic excitation from cortical interneurons, from simultaneous inputs dispersed along the dendritic tree of corticospinal output cells [10, 27] or from mechanism involving back-propagating action potentials coinciding with subthreshold distal EPSP causing pyramidal cells to respond by discharging a burst of action potentials [11, 28]. Regardless of the underlying mechanisms, the I-wave periodicity reflects a time windows of heightened excitability, during which a consecutive TMS pulse more readily excites corticospinal output neurons [22]. The observed multi-pulse facilitation at the first SICF peak IPI in the present work is in good agreement with previous results.

Importantly, the tenet that SICF and I-wave periodicity are underpinned by the same neuronal circuitry remains to be proven. It is possible that the facilitatory effects arise because the first suprathreshold stimulus renders initial axonal segments of neurons in the late I-wave pathway hyperexcitable and thereby increase the sensitivity of these axonal structures to later stimuli [29]. The period of increased sensitivity starts after refractory and is curtailed by the small timeconstant characteristic of the membrane at the initial segment. This gives a ‘facilitation window’ around 1.5 ms corresponding to the peak1 IPI. While the notion of a pre-pulse sensitization provides a potential mechanistic route for the observed MEP facilitation with multi-pulse TMS adjusted to the first SICF peak, it fails to explain the occurrence of later SICF peaks. Likewise, the demonstration of paired-pulse facilitation at peak1-latency and peak2-latency in our second follow-up experiment supports a model that attributes this short-latency facilitation to I-wave interactions at the cortical level.

### Short-latency facilitation by multi-pulse TMS independently of I-wave periodicity

The facilitatory effect of multi-pulse TMS was not restricted to the temporal window set by I-wave periodicity, because multi-pulse TMS_HAND_ at trough1-latency also resulted in corticomotor facilitation albeit to a lesser extent than multi-pulse TMS_HAND_ at peak1-latency. TMS_HAND_ can produce a peripheral motor response by exciting cervical motoneurons via low-threshold, direct (monosynaptic), corticomotoneuronal projections and high-threshold, indirect (disynaptic), corticomotoneuronal projections to segmental interneurons [30, 31]. Compared to multi-pulse TMS_HAND_ at peak1-latency, I-wave independent facilitation became gradually more pronounced with stimulation intensity. Indeed, the stimulus-response curves obtained with multi-pulse TMS_HAND_ at trough1-latency and peak1-latency TMS converged at high stimulus intensities (figure 3). We attribute the gradual emergence of I-wave independent facilitation with increasing stimulus intensity to a more efficient recruitment of high-threshold polysynaptic projections from both rostral and caudal M1 as described by Witham et al. (2016). Additionally, multi-pulse facilitation at higher stimulus intensities may be caused by summation of transsynaptic excitation in other descending pathways, such as propriospinal [31–33]; reticulospinal[34, 35] or the convergence of these on intercalated interneurons [36].

Regardless of whether the I-wave independent facilitation is due to temporal summation in descending pathways or in circuitries upstream to the output neurons of M1, the overall summation reflected in MEP facilitation was less efficient for multi-pulse TMS_HAND_ at trough1 latency than peak1 latency.

### Effects of number of pulses on excitation of the corticospinal output

We systematically varied the number of stimuli and found a consistent, albeit moderate, effect of the number of pulses on the magnitude of short-latency corticomotor facilitation. In the main experiment, the overall facilitatory effect of the number of pulses was significant in full factorial ANOVAs on normalized multi-pulse facilitation. This observation was replicated in a follow-up experiment, in which we probed short-latency corticomotor facilitation at sub-threshold stimulus intensities. The corticomotor response increased with the number of pulses. Together, our results show that there is a moderate but consistent facilitatory effect on MEP facilitation with increasing number of pulses independent of I-wave periodicity.

Our finding of I-wave independent additional facilitation from adding a third or fourth pulse to the paired pulses contrasts recent findings. Sacco et al. [22] found a strong extra-facilitatory effect of the third pulse in seven individuals when applying triple-pulse TMS_HAND_, but only at I-wave periodicity. The discrepancy could be attributed to differences in stimulation parameters. Sacco et al. used monophasic TMS pulses producing a P-A directed current in the precentral gyrus. Although the second effective phase of the biphasic pulse delivered in the present study also produced P-A directed currents, it cannot be excluded that the mechanisms of action may differ slightly between pulse shapes. Noteworthy, another triple-pulse TMS_HAND_ study demonstrated complex multi-pulse facilitatory effects, which the authors attributed to interactions between intracortical circuits causing SICF and short-latency intracortical inhibition (SICI) i.e., the reduction in MEP size observed when conditioning a suprathreshold pulse by subthreshold pulse with IPIs below 4.5ms [37]. It is possible that in our study, additional I-wave dependent MEP facilitation by the third and fourth pulses was counteracted by concurrent TMS-induced intracortical inhibition. A similar phenomenon could occur at the level of the spinal cord. The IPI between the first and third or fourth pulse is sufficient to render the effects of the latter sensitive to di- or polysynaptic inhibition through Ia interneurons [38].

### Strengths and limitations

This study has several strengths: we concurrently assessed the facilitatory effects of varying TMS bursts covering a large range of TMS intensities. Furthermore, we investigated I-wave periodicity with high sensitivity through personalization of the IPI based on the individual SICF latencies.

The study also has limitations: first, we used a biphasic pulse shape and did not systematically study the impact of different TMS-induced current directions. In contrast to the majority of SICF (e.g. [16, 17]), we chose a biphasic pulse shape and the same stimulation intensity throughout each multi-pulse burst. Notwithstanding, peak latencies and periodicity of SICF were very similar in our study compared to a seminal monophasic SICF study [16], with a small delay which is in accordance with the findings by Di Lazzaro et al. [39]. In line with previous findings [40], the latencies and relative facilitation of biphasic SICF were comparable with the those revealed by monophasic SICF, thereby confirming, that SICF can be probed with biphasic TMS pulses. Importantly, all participants displayed two clear SICF peaks separated by a trough, but with substantial interindividual variability of the IPIs, underlining the importance of individualization when targeting I-wave interactions. Furthermore, the use of biphasic pulses also enables the results to be readily applied in future rTMS studies. Second, since all experiments were performed in the absence of muscle activity, caution is wanted when comparing the present findings to the effects of pulse-number with IPIs delivered during a background contraction.

## Conclusions

To the best of our knowledge, our study is the first to show two distinct mechanisms that contribute to the facilitatory effects of multi-pulse TMS_HAND_ at ultra-high repetition rates. In addition to the well-known facilitatory effect related to I-wave periodicity, we found evidence for multi-pulse facilitation at an inter-pulse interval that is unrelated to I-wave periodicity. The extent to which MEP facilitation following repetitive multi-pulse TMS_HAND_ can be ascribed to mechanisms dependent versus independent of I-wave periodicity remains to be explored. Our results are of relevance for the future use of repetitive multi-pulse TMS_HAND_ as a means of inducing corticospinal plasticity in the human motor system and underline that an exclusive focus on I-wave periodicity may be too narrow, when aiming at understanding how TMS excites the precentral motor representations and induces plasticity.

**Table 1:**
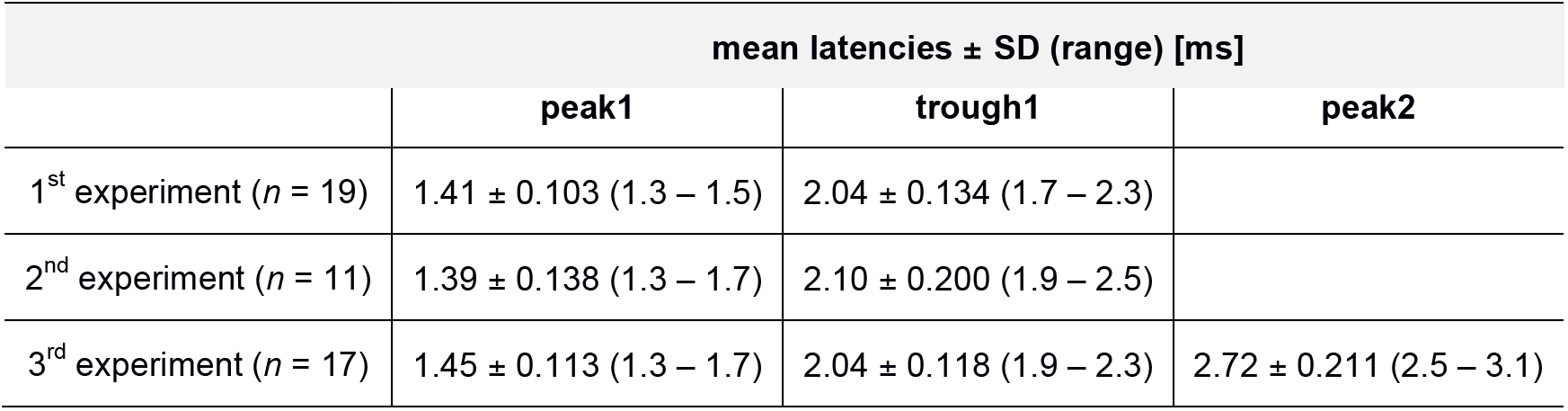
Obtained peak- and trough-latencies using SICF for all experimental sessions

## Additional information

### Data Availability Statement

All data that support the findings of this study are openly available in “Open Science Framework” at https://osf.io/acges/ (DOI 10.17605/OSF.IO/ACGES).

## Acknowledgements

This work is part of the project “Biophysically adjusted state-informed cortex stimulation” (BASICS) funded by a synergy grant from Novo Nordisk Foundation (Interdisciplinary Synergy Program 2014; grant number NNF14OC0011413)

